# Stress Granule Protein TZF1 Enhances Salt Stress Tolerance by Targeting *ACA11* mRNA for Degradation in Arabidopsis

**DOI:** 10.1101/2024.01.12.575397

**Authors:** Siou-Luan He, Bin Li, Walter J. Zahurancik, Henry C. Arthur, Vaishnavi Sidharthan, Venkat Gopalan, Lei Wang, Jyan-Chyun Jang

**Affiliations:** Department of Horticulture and Crop Science, The Ohio State University, Columbus, OH, USA; Center for Applied Plant Sciences, The Ohio State University, Columbus, OH, USA; Center for RNA Biology, The Ohio State University, Columbus, OH, USA; China Key Laboratory of Plant Molecular Physiology, Institute of Botany, Chinese Academy of Sciences, and University of Chinese Academy of Sciences, Beijing, China; Department of Chemistry and Biochemistry, The Ohio State University, Columbus, OH, USA; Academician Workstation of Agricultural High-Tech Industrial Area of the Yellow River Delta, National Center of Technology Innovation for Comprehensive Utilization of Saline-Alkali Land, Shandong, China

## Abstract

Tandem CCCH zinc finger (TZF) proteins play diverse roles in plant growth and stress response. Although as many as 11 TZF proteins have been identified in *Arabidopsis*, little is known about the mechanism by which TZF proteins select and regulate the target mRNAs. Here, we report that *Arabidopsis* TZF1 is a bona-fide stress granule protein. Ectopic expression of *TZF1* (*TZF1 OE*), but not an mRNA binding-defective mutant (*TZF1^H186Y^ OE*), enhances salt stress tolerance in *Arabidopsis*. RNA-seq analyses of NaCl-treated plants revealed that the down-regulated genes in *TZF1 OE* plants are enriched for functions in salt and oxidative stress responses. Because many of these down-regulated mRNAs contain AU- and/or U-rich elements (AREs and/or UREs) in their 3’-UTRs, we hypothesized that TZF1—ARE/URE interaction might contribute to the observed gene expression changes. Results from RNA immunoprecipitation-quantitative PCR analysis, gel-shift, and mRNA half-life assays indicate that TZF1 binds and triggers degradation of the *autoinhibited Ca^2+^-ATPase 11* (*ACA11*) mRNA, encoding a tonoplast-localized calcium pump that extrudes calcium and dampens the signal transduction pathways necessary for salt stress tolerance. Furthermore, this salt stress-tolerance phenotype was recapitulated in *aca11* null mutants. Remarkably, a set of positive regulators for salt stress tolerance was upregulated in *TZF1 OE* plants. These include *Na^+^/H^+^ Exchanger* (*NHX*) family members known to contribute to Na^+^ homeostasis and salinity stress tolerance. Collectively, we present a model in which TZF1 targets *ACA11* and *ACA4* directly, and repressors of *NHXs* and other negative regulators indirectly for mRNA degradation to enhance plant salt stress tolerance.

## INTRODUCTION

Salinity stress is harmful to most non-halophytes plants, seriously limiting crop growth and productivity (Zhao et al., 2021). Plants have developed sophisticated mechanisms to temporarily adapt to high salt environments by altering gene expression, physiology, and metabolism (Zhu, 2002; Deinlein et al., 2014; Julkowska and Testerink, 2015; Zhu, 2016). Thus, unraveling the genetic mechanisms underlying salt stress tolerance could provide important clues to improve crop fitness and yields under prolonged salinity stress or in the regions with high soil salinity.

*Arabidopsis* tandem CCCH zinc finger (AtTZF) proteins play a key role in regulating various plant hormone-mediated growth and environmental responses. There are 11 *TZF* genes in *Arabidopsis* that are differentially expressed temporally and spatially (Jang, 2016). In the *AtTZF* gene family (hereafter as *TZF*), *TZF1* is ubiquitously expressed at a high level and the best characterized family member. TZF1 is a positive regulator of abscisic acid (ABA), sugar, and salt stress tolerance responses, while serving as a negative regulator of gibberellin (GA) responses (Lin et al., 2011; Han et al., 2014). In rice, OsTZF1, the homologous gene of *TZF1*, is involved in seed germination, seedling growth, leaf senescence, and oxidative stress tolerance (Jan et al., 2013). TZF2 and TZF3 act as positive regulators for ABA, oxidative, and salt stress responses, and as negative regulators for stress hormone methyl jasmonate (MeJA)-induced senescence (Huang et al., 2011; Huang et al., 2012; Lee et al., 2012). Furthermore, rice OsTZF2 (OsDOS), the homolog of TZF2, can delay MeJA-induced leaf senescence (Kong et al., 2006). TZF4/5/6 are positive regulators of ABA responses, while acting as negative regulators for GA and phytochrome-mediated seed germination responses (Kim et al., 2008; Bogamuwa and Jang, 2013). TZF7 to TZF 11 are positive regulators of vegetative growth and abiotic and biotic stress tolerance responses, while functioning as negative regulators for stress-induced transition to flowering (Sun et al., 2007; Blanvillain et al., 2011; Maldonado-Bonilla et al., 2014; Kong et al., 2021). Although TZF1/2/3/10/11 proteins have been shown to be redundantly involved in salt stress responses (Jang, 2016), the molecular mechanisms by which TZF proteins target mRNAs to govern the salinity stress response are still unclear.

Ribinucleoprotein (RNP) granules are membrane-less biomolecular condensates generally form via liquid-liquid phase separation mediated by multivalent protein-protein, protein-RNA, and RNA-RNA interactions. The scaffold proteins in the RNP granules are characterized by having intrinsically disordered, low-complexity, or prion-like domains (Ripin and Parker, 2023; Solis-Miranda et al., 2023). Two of the best characterized cytoplasmic RNP granules are processing bodies (PBs) and stress granules (SGs), which are dynamically assembled in response to environmental stresses (Jang et al., 2020). Human tristetraprolin (hTTP), an intensively-studied prototypical TZF protein, controls mRNA stability by selective binding to the target genes (Carballo et al., 1998). hTTP often binds AU-rich elements (AREs) in the 3’-UTR of mRNA and recruits target mRNAs to PBs and SGs for gene silencing through RNA decay and translational repression (Lai et al., 1999). In *Arabidopsis*, TZF1 was found to traffic between cytoplasmic foci and nuclei and was colocalized with PB (DCP2) and SG (PABP8) markers (Pomeranz et al., 2010). TZF1 could also bind polyU (Pomeranz et al., 2010) and ARE (Qu et al., 2014) with specificity. Furthermore, TZF1 could directly bind 3’-UTR of *Target of Rapamycin* (*TOR*) mRNA using its tandem zinc finger motif to affect *TOR* mRNA stability. The interaction of TZF1-*TOR* mRNA is necessary for root meristem cell proliferation by integrating both transcriptional and post-transcriptional regulation of gene expression (Li et al., 2019). Although the molecular mechanisms of hTTP in promoting mRNA degradation in animals has been characterized in great detail (Brooks and Blackshear, 2013), the mechanisms by which plant TZF proteins bind and elicit degradation of target mRNAs are poorly understood.

Plant TZF proteins play critical roles in salt, drought, oxidative, and many other stress responses, but the link between TZF1-mediated stress tolerance responses and TZF1-mediated mRNA degradation has not been well-established. To date, it remains a challenge to unbiasedly identify genome-wide TZF1 target mRNAs and specific binding sites in vivo. Here, we show that TZF1 could enhance salt stress tolerance by binding to and initiating degradation of mRNAs of salt- or oxidative-stress responsive genes. As proof of principle, we performed detailed biochemical and molecular analyses to demonstrate that TZF1 enhances salt stress tolerance by binding to a specific 3’-UTR region of a down-regulated target gene *ACA11*, a negative regulator in salt stress signaling.

## RESULTS

### TZF1 Is Involved in Salt Stress Tolerance

Previous studies have shown that *AtTZF1* transcription was upregulated by high salt and that *TZF1* overexpression plants enhanced salt stress tolerance through regulating cellular ion balance and limiting oxidative and osmotic stress (Han et al., 2014). To further investigate the underlying mechanism of TZF1 in enhancing plant salt stress tolerance and whether the RNA-binding ability of TZF1 is required for enhancing salt stress tolerance, *Arabidopsis* plants overexpressing wild-type TZF1 (*TZF1 OE*) (Lin et al., 2011) and TZF1 mutant containing a histidine to tyrosine substitution in its second zinc finger motif (*TZF1^H186Y^ OE*) were employed in this study (Li et al., 2019). The *TZF1^H186Y^ OE* mutant allele was obtained from a genetic screen of ethyl methanesulfonate (EMS)-mutagenized *TZF1 OE* homozygous population and was isolated as a *TZF1 OE* intragenic mutant of the transgene *CaMV35S:TZF1-GFP*. Notably, *TZF1 OE* plants are dwarf and late flowering while *TZF1^H186Y^ OE* plants are morphologically similar to the wild-type (Col-0) plants with normal flowering time. The reversion of the overexpression phenotypes is likely due to loss-of-function of mutated transgene *CaMV35S:TZF1^H186Y^-GFP*, as evidenced by a previous study (Li et al., 2019).

Wild-type, *TZF1 OE*, *TZF1^H186Y^ OE*, and *tzf1* T-DNA knockout seedlings were transferred to MS medium with or without NaCl or sorbitol treatment. There were no observable differences in phenotypes between wild-type, *tzf1*, *TZF1 OE* and *TZF1^H186Y^ OE* seedlings in the absence of NaCl treatment (Fig. 1A; Supplemental Fig. S1). However, under salt stress (200 mM NaCl), the survival rate, fresh weight, and chlorophyll content of the wild-type, *tzf1*, and *TZF1^H186Y^ OE* seedlings were significantly reduced, and most seedling leaves showed bleached phenotypes. In contrast, the survival rate and chlorophyll content of *TZF1 OE* were significantly higher than that of the rest of the plants (Figs. 1A-D; Supplemental Fig. S1). Noticeably, no significant differences on the leaf bleach phenotypes were observed among plants treated with 300 mM or 400 mM sorbitol (Supplemental Fig. S2), indicating that TZF1 is more likely to be specifically involved in salt stress response, but not general osmotic stress response. To further confirm that *TZF1^H186Y^ OE* conferred a revertant phenotype of *TZF1 OE*, we recapitulated the mutagenesis event by overexpressing *TZF1^H186Y^* directly in the wild-type background under the control of the *CaMV 35S* promoter. Upon treatment with 200 mM NaCl, *TZF1^H186Y^ OE L8* seedlings exhibited a high mortality rate (Supplemental Fig. S3) as was found in the original EMS allele of *TZF1^H186Y^ OE*.

**Figure 1.**
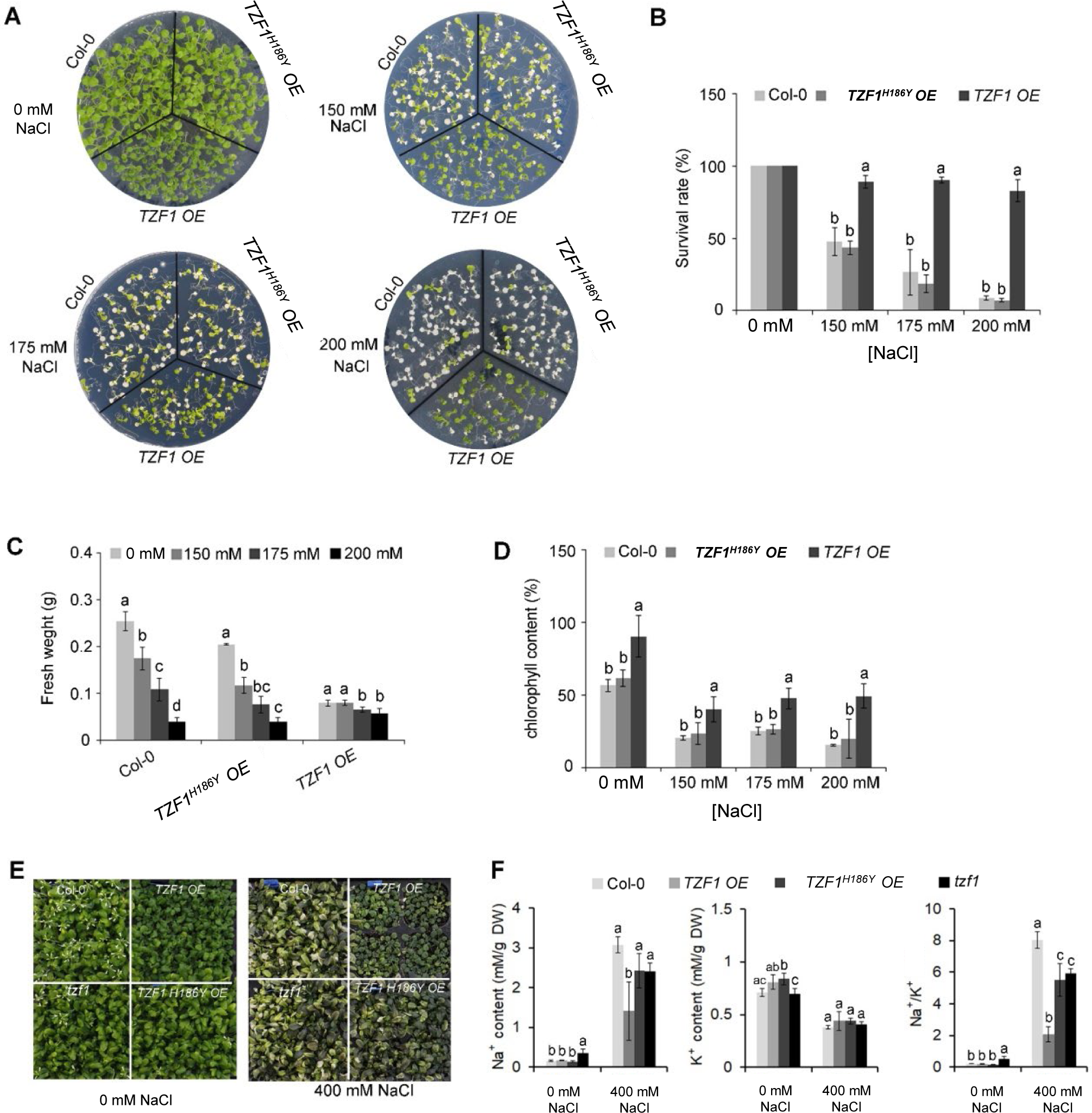
Overexpression of *AtTZF1* improves salt stress tolerance. A, Phenotypes of the Col-0, *TZF1 OE*, and *TZF1^H186Y^ OE* plants under normal and salt stress conditions. Seven-day-old seedlings grown under long-day condition (16/8 h light/dark) were transferred to MS plates containing different NaCl concentrations for additional eight days. B, Survival rates of seedlings shown in (A). C, Fresh weights of seedlings shown in (A). D, Chlorophyll contents of seedlings shown in (A). E and F, Effects of NaCl treatment on Na^+^ and K^+^ contents, and Na^+^/K^+^ ratios in various plants. Eighteen-day-old plants grown in 12 h light/dark cycles were treated with 400 mM NaCl or water (0 mM NaCl) every three days for two weeks. Data represent the average of three replicates ± *SD*. Different letters (*a*, *b*, and *c*) indicate significant differences at *P*< 0.05 by one-way ANOVA analysis using the SPSS software.

Under salt stress, plants utilize specific mechanisms to modulate Na^+^/K^+^ homeostasis. Thus, a low cytosolic Na^+^/K^+^ ratio is a key indication of salt stress tolerance (Zhao et al., 2021). Na^+^ and K^+^ concentrations and Na^+^/K^+^ ratio were measured in leaves of *Arabidopsis* plants subjected to 400 mM NaCl in soil for 14 days. Under such treatment, growth of the wild-type, *tzf1*, and *TZF1^H186Y^ OE* plants was inhibited, while *TZF1 OE* had fewer withered yellow leaves compared to the other plants (Fig. 1E). Moreover, *TZF1 OE* plants showed a reduced accumulation of Na^+^ ions and exhibited a lower intracellular Na^+^/K^+^ ratio during salt stress compared to wild-type plants (Fig. 1F). These results indicate that *TZF1* overexpression results in high salt-stress tolerance, suggesting that TZF1 functions as a positive regulator of salt-stress tolerance in *Arabidopsis*.

### *TZF1* Overexpression Alleviated the Oxidative Damages Caused by Salt Stress

The production of hydrogen peroxide (H_2_O_2_), a major reactive oxygen species (ROS) induced by salt stress, is an indicator of oxidative damage. To further assess the role of TZF1 in salt stress response, *TZF1 OE*, *TZF1^H186Y^ OE*, and wild-type seedlings and adult plant leaves were stained with 3,3’-diaminobenzidine (DAB) to determine the H_2_O_2_ levels under normal and stress conditions. Under normal growth conditions, the DAB staining of all plant leaves showed no differences. However, in the presence of 175 mM NaCl, the staining intensity of the *TZF1 OE* plants was significantly lower than that of the wild-type and *TZF1^H186Y^ OE* plants (Fig. 2). These results indicate that H_2_O_2_ accumulation in the wild-type and *TZF1^H186Y^ OE* plant leaves was greater than that in the *TZF1 OE* plants during salt stress and suggest that *TZF1* overexpression enhanced salinity tolerance by reducing ROS accumulation.

**Figure 2.**
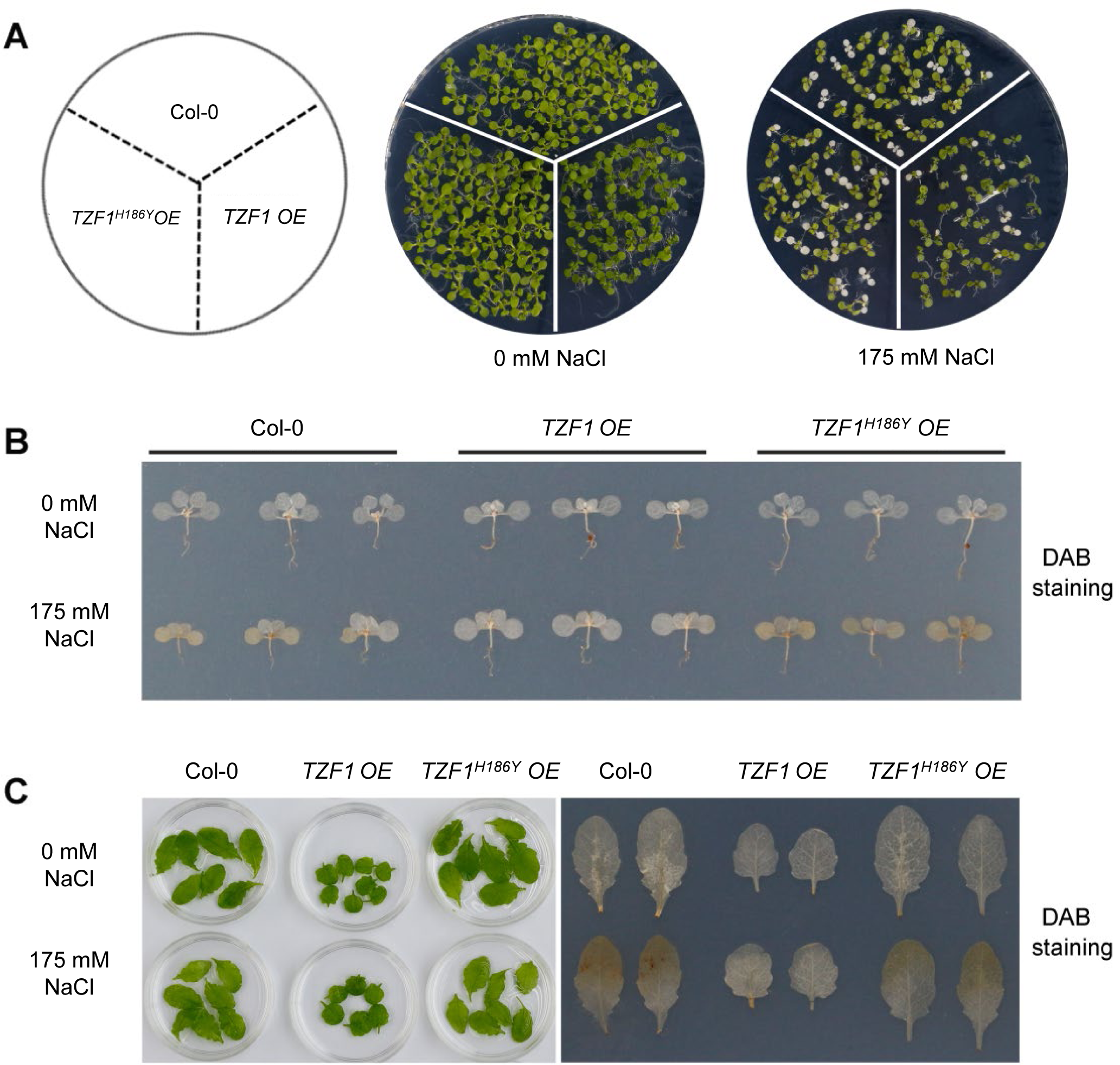
*TZF1 OE* plants accumulate less H_2_O_2_ under salt stress. A, Phenotypes of the Col-0, *TZF1 OE*, and *TZF1^H186Y^ OE* under normal and stress conditions. Seven-day-old seedlings grown under long-day condition were transferred to MS plates containing 175 mM NaCl for another 5 days. B, DAB staining to determine the levels of H_2_O_2_ accumulation in various seedlings shown in (A). C, DAB staining to determine the H_2_O_2_ accumulation levels in the detached leaves of eighteen-day-old plants grown in soil under 12 h light/dark cycles treated with 175 mM NaCl or water (0 mM NaCl) for additional three days.

### Salt-Induced TZF1 Stress Granule Assembly

Previous studies showed that TZF1 could colocalize with both PB and SG markers in *Arabidopsis* protoplasts. TZF1 localized to cytoplasmic foci in intact plants and these cytoplasmic granules can be induced by either wounding or MeJA treatment (Pomeranz et al., 2010). Moreover, the rice OsTZF1 cytoplasmic foci could be induced by NaCl treatment (Jan et al., 2013). To investigate if AtTZF1 cytoplasmic foci could be induced by high salt as well, we examined seven-day-old seedlings of *TZF1 OE* and *TZF1^H186Y^ OE* root cells with or without 200 mM NaCl treatment at 4 h and 24 h. Results showed that NaCl treatment could induce the formation of cytoplasmic foci in both *TZF1 OE* and *TZF1^H186Y^ OE* plants (Fig. 3). Interestingly, salt-induced TZF1-GFP foci appeared to be more abundant at 24 h in *TZF1^H186Y^ OE* plants than *TZF1 OE* plants, suggesting that TZF1^H186Y^-GFP protein might be more stable, albeit non-functional in salt stress tolerance. As the green auto-fluorescence from the cytoplasm of the light-grown seedlings interfered with the observation of GFP foci, etiolated seedlings were further examined. Again, the results confirmed that TZF1-GFP foci were salt-inducible. Consistently, both diffused GFP and cytoplasmic granule signals were stronger in *TZF1^H186Y^ OE* plants than that in *TZF1 OE* plants before and after NaCl treatment (Supplemental Fig. S4).

**Figure 3.**
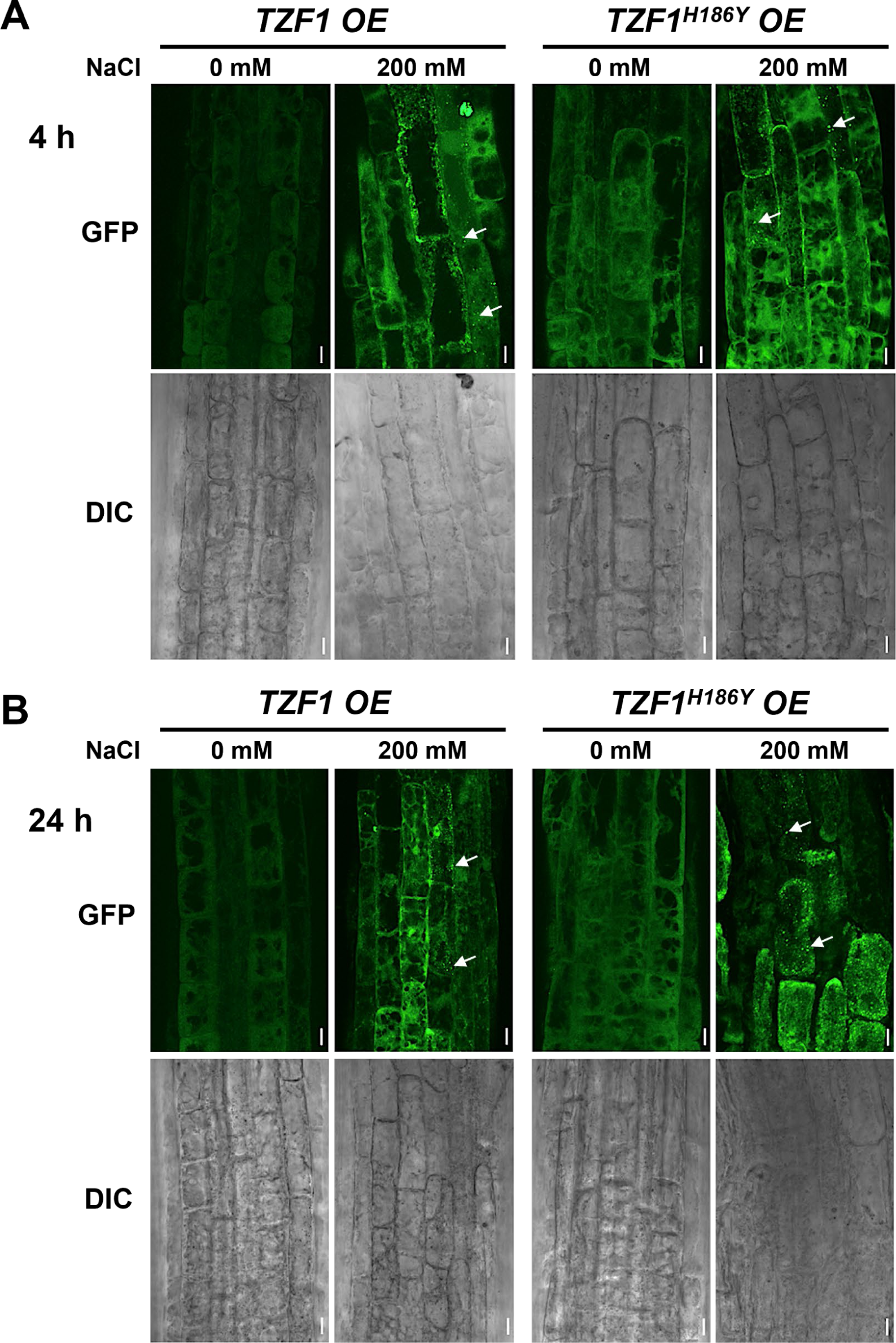
TZF1 cytoplasmic foci induced by salt stress. A and B, Confocal microscopy of root cells of seven-day-old *TZF1 OE* and *TZF1^H186Y^ OE* seedlings treated with 200 mM NaCl for 4 h (A) and 24 h (B), respectively. GFP signals were generated by the expression of *CaMV35S:TZF1-GFP* and *CaMV35S:TZF1^H186Y^-GFP* fusion genes. Typical TZF1 cytoplasmic foci are indicated by arrows. Salt-induced cytoplasmic foci appeared to be more abundant in *TZF1^H186Y^ OE* plants, particularly after 24 h treatment. Scale bars = 10 μm.

Although OsTZF1 cytoplasmic foci were induced by salt stress, the identities of the foci were unknown (Jan et al., 2013). We have shown previously that AtTZF1 could completely co-localize with both PB (DCP2) and SG (PABP8) markers (Pomeranz et al., 2010). It was therefore confusing whether TZF proteins are components of PBs, SGs, or both. Subsequently, it was found that although plant DCP2 is a major component of mRNA decapping complex, it was not a PB-specific marker (Motomura et al., 2014), raising a possibility that TZF1 might be mainly localized in SGs and could be recruited or exchanged to PBs under specific cues. To test this hypothesis, AtTZF1 sub-cellular localization was re-examined using a set of different markers in *Arabidopsis* protoplast transient expression analyses (Sheen, 2001). Results showed that AtTZF1 could not or only very partially co-localize with authentic PB markers DCP1 and DCP5, respectively, but completely co-localize with SG markers G3BP and UBP1b (Sorenson and Bailey-Serres, 2014; Jang et al., 2020; Solis-Miranda et al., 2023). Importantly, both TZF1-GFP and TZF1^H186Y^-GFP were completely co-localized with SG marker UBP1b (Fig. 4A). To determine if the difference in protein abundance in intact plants was specific to fusion proteins with GFP tag, *TZF1-mCherry* and *TZF1^H186Y^-mCherry* were independently expressed in *Arabidopsis* protoplast. Results showed that the signal of TZF1^H186Y^-mCherry was still much stronger than TZF1-mCherry (Fig. 4B), suggesting that the former protein might be more stable hence accumulated at higher level.

**Figure 4.**
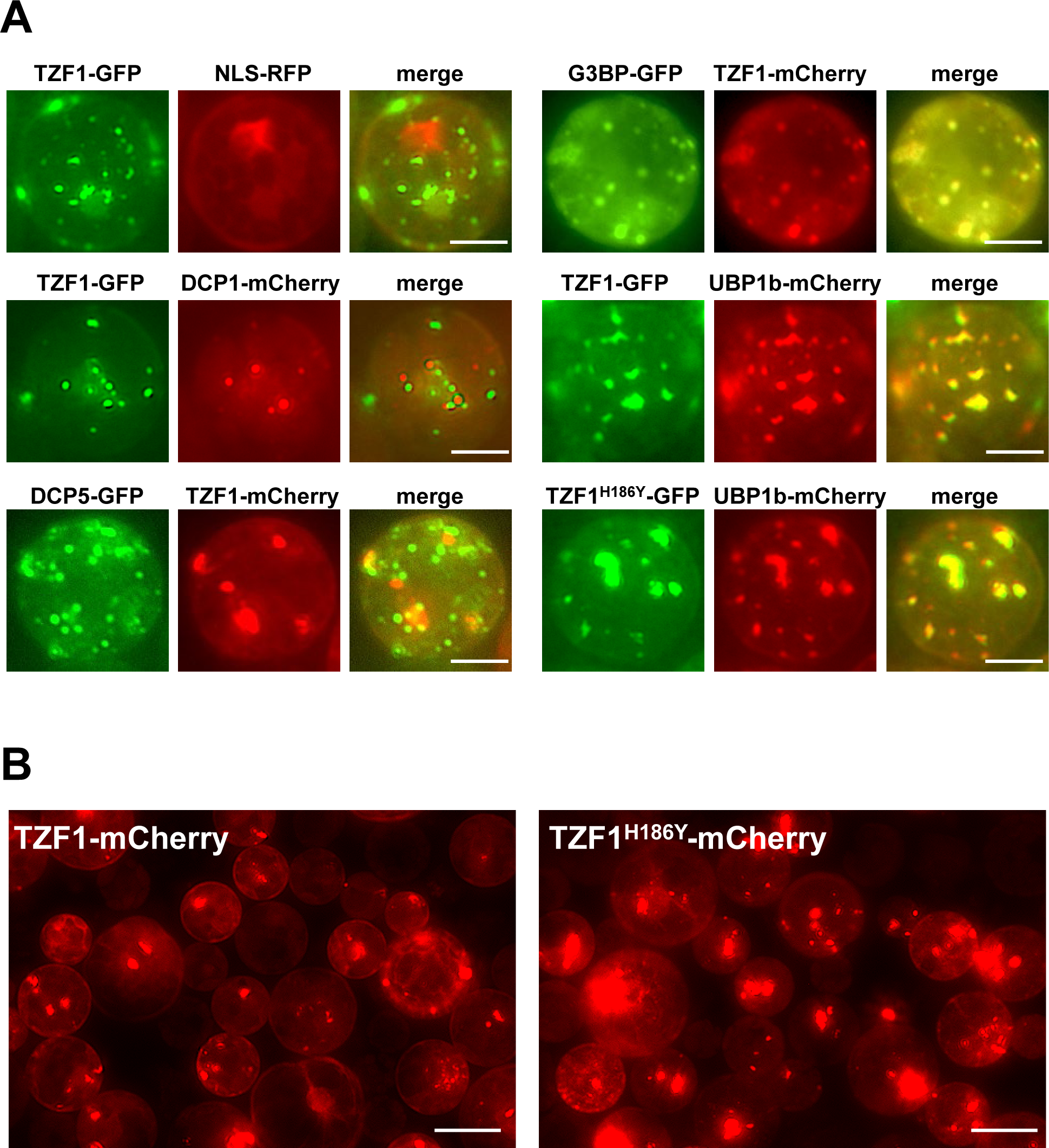
TZF1 is localized in SGs. A, Individual pair of reporting constructs were co-expressed in *Arabidopsis* protoplasts. TZF1 was not colocalized with P-body markers DCP1, very partially co-localized with DCP5, but completely co-localized with SG markers G3BP or UBP1b. Scale bars = 10 μm. B, Cytoplasmic granules were more abundant in cells expressing TZF1^H186Y^-mCherry than TZF1-mCherry. Scale bars = 20 μm.

To further determine if the salt-induction of TZF1 SG assembly was a result of change in protein abundance, a time-course immunoblot analysis using seven-day-old seedlings was conducted. Results showed that the protein level of either TZF1-GFP or TZF1^H186Y^-GFP was unaffected by salt treatment and remained nearly constant during the time course (Fig. 5A). Therefore, salt induced TZF1 SGs assembly was mediated by unknown post-translational regulatory mechanisms. Noticeably, TZF1^H186Y^-GFP was accumulated at a higher level than TZF1-GFP before and after salt treatment (Fig. 5A). To determine protein stability, time-course analysis using seven-day-old seedlings treated with protein synthesis inhibitor cycloheximide (CHX), proteosome inhibitor MG115/132, and a combination of CHX and MG115/132 was conducted. Results showed that TZF1 protein was extremely unstable—it almost completely disappeared after being treated by CHX for just 1 h (Fig. 5B). By contrast, its accumulation was enhanced by MG115/132. Additionally, TZF1 protein at 1 h mark had already been degraded nearly 50%, suggesting that the slow action of MG132/115 treatment could not prevent the fast turnover of TZF1. This was evidenced by the combined CHX and MG115/132 treatment in which TZF1 had already been degraded at 1 h mark before MG115/132 could protect it from being degraded. Compared to TZF1-GFP, TZF1^H186Y^-GFP was remarkably stable (Fig. 5B), confirming that the higher accumulation of steady-state TZF1^H186Y^-GFP or TZF1^H186Y^-mCherry was due to enhanced protein stability. Together, these results suggested that although TZF1^H186Y^-GFP is more stable and accumulated at higher levels, it is non-functional in enhancing plant salt stress tolerance, albeit it could still localize to SGs.

**Figure 5.**
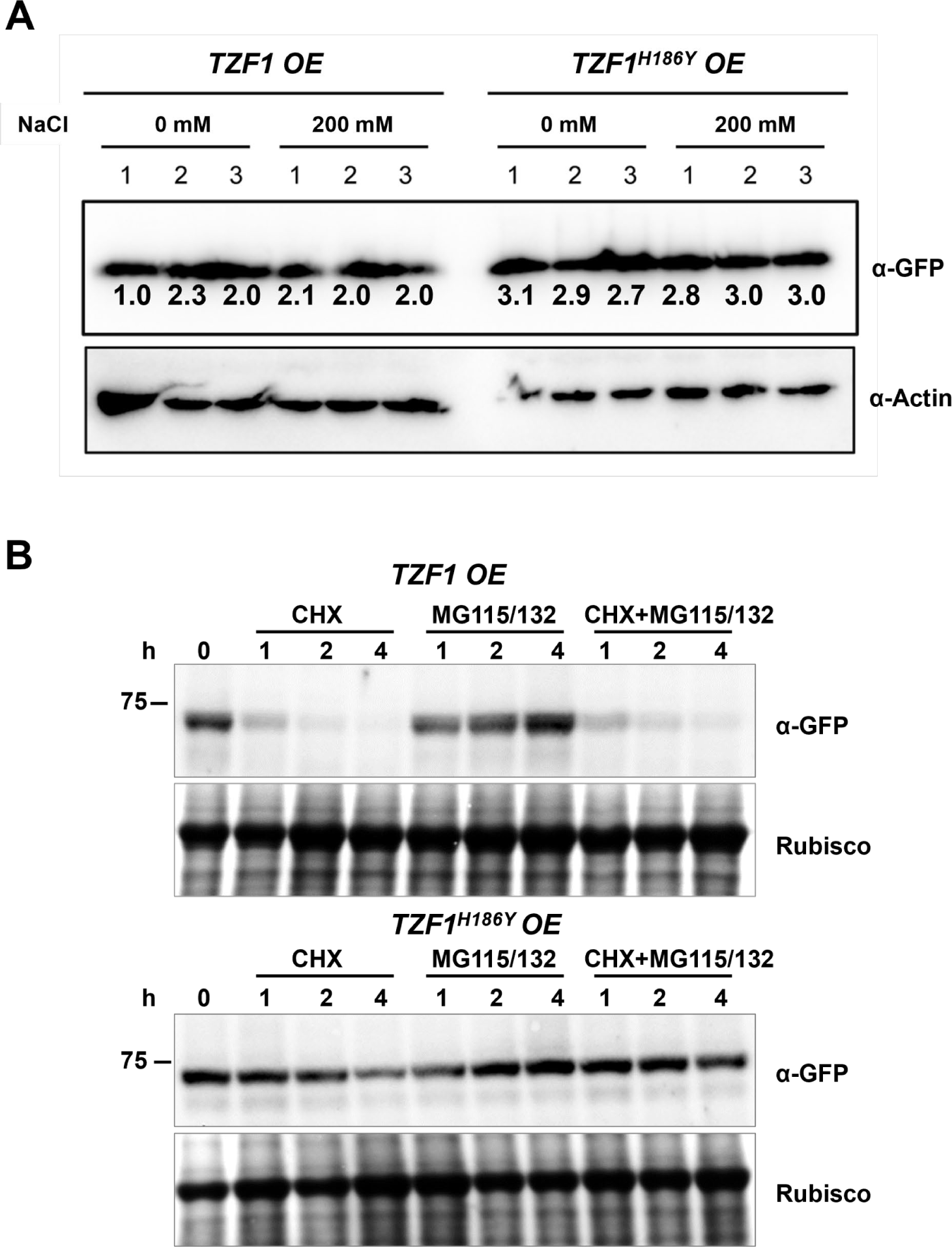
TZF1-GFP is less stable than TZF1^H186Y^-GFP. A, Seven-day-old seedlings were treated with 150 mM NaCl in a time-course experiment. Samples were collected at indicated times and total proteins were extracted for immunoblot analysis. The number below each protein band is normalized value of GFP vs ACTIN signal. Note that the low value of the first sample is due to the smear of the ACTIN band. Three independent samples were examined for each treatment. B, Half-life analysis to determine protein stability. Seven-day-old seedlings were incubated with 30 μM CHX, 50 μM MG115/132, or both CHX and MG115/132. Seedlings were collected at different time points, and then total proteins were extracted for immunoblot analysis. Rubisco was used as a loading control.

### TZF1 Affects Salt-Induced Transcriptome Change

To uncover the role of TZF1 in modulating transcript levels of salt stress-related genes, we performed deep sequencing of RNA extracted from the wild-type, *TZF1 OE*, and *TZF1^H186Y^ OE* plants treated with 150 mM NaCl. In total, there were nearly 1,000 up-regulated and 1176 down-regulated differentially expressed genes (DEGs) present in the *TZF1 OE* plants (Fig. 6A; Supplemental Fig. S5A). Gene ontology (GO) analysis based on biological process clustering of DEGs in *TZF1 OE* plants revealed that up-regulated DEGs are over-represented by genes that are subject to ubiquitin transferase activity (Supplemental Fig. S5B) while down-regulated DEGs preferentially associated with response to stress and response to stimulus (Fig. 6B). Although the analysis of down-regulated DEGs in *TZF1 OE* is the focus of this report, we also highlight below the nexus between the outstanding up-regulated DEGs (Supplemental Fig. S5C) and salt stress tolerance (see *Discussion*). For example, Na^+^/H^+^ antiporters (NHXs) (Bassil et al., 2011), CBL-interacting protein kinase 8 (CIPK8) (Yin et al., 2020), RGA-LIKE3 (RGL3) (Shi et al., 2017), SCF E3 ligase (PP2-B11) (Jia et al., 2015), annexin (AnnAt1) (Jia et al., 2015), and vacuolar protein sorting 23A (VPS23A) (Lou et al., 2020) are all implicated as positive regulators in salt stress tolerance response.

**Figure 6.**
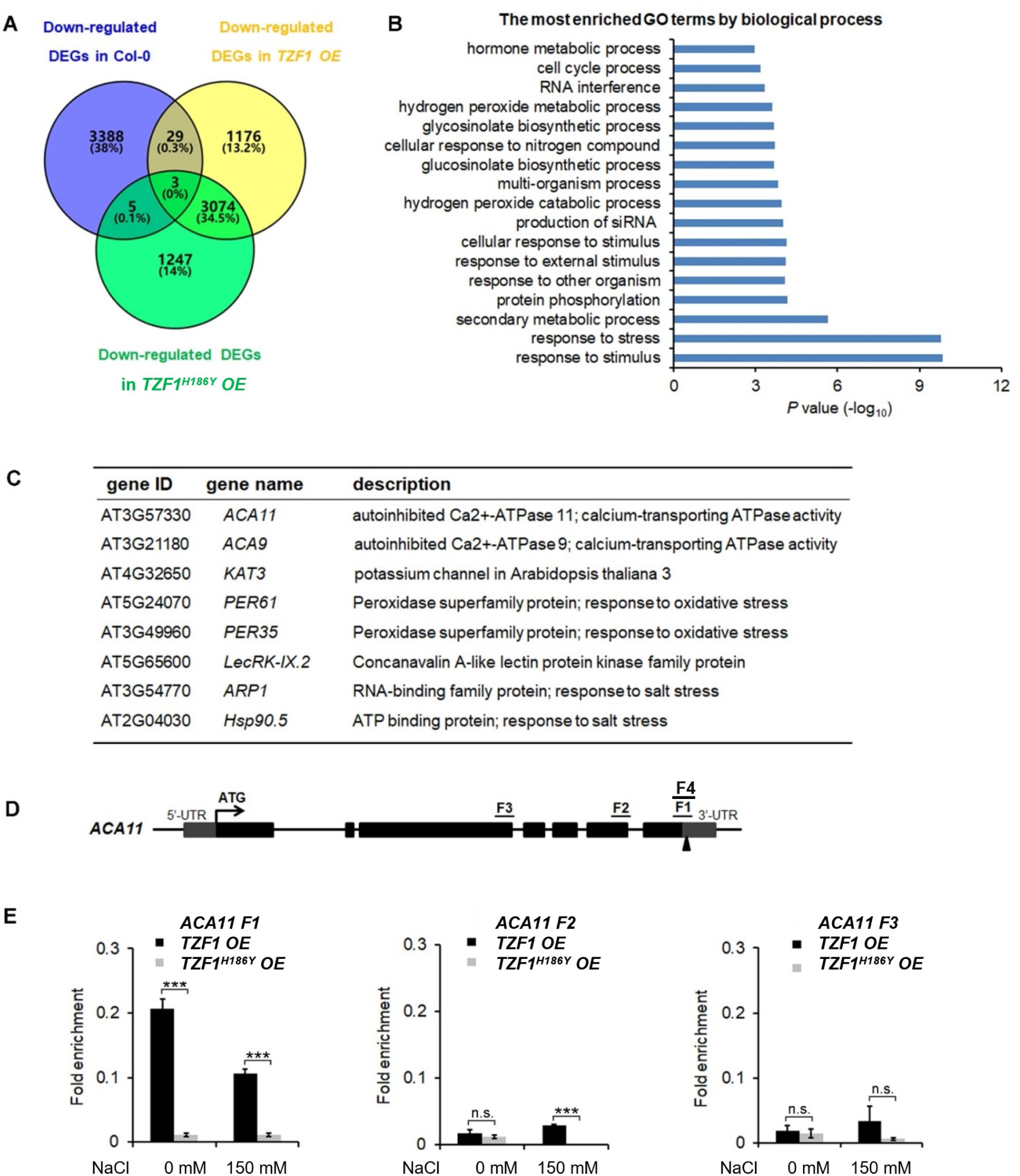
TZF1 affects salt stress responsive genes. A, Venn diagram analysis of down-regulated DEGs in the Col-0, *TZF1 OE*, and *TZF1^H186Y^ OE* plants treated with NaCl. RNA-seq analysis was conducted using eleven-day-old seedlings grown on MS plates under 12 h light/dark cycles and treated with 150 mM NaCl for 4 h. B, GO analysis of DEGs specifically found in NaCl-treated *TZF1 OE* plants. *P* values were generated by Fisher’s exact test. C, A set of the salt stress or oxidative stress related genes that was down-regulated in *TZF1 OE* plants and contained AU- or U-rich elements in the 3’-UTRs was selected for further analysis. D, Diagram depicting the probe regions (F1 to F3) on *ACA11* gene used for RIP-qPCR analysis. Arrowhead indicates an ARE containing region. E, RIP-qPCR results indicated a potential binding of TZF1 to F1 region of the *ACA11* mRNA. Data represent the average of 3 replicates ± SD. Asterisks indicate significant differences (***, *P* < 0.001) by Student’s *t* test.

It has been shown that the TZF motif of TZF1 is required for both RNA targeting and turnover (Qu et al., 2014). Additionally, TZF1^H186Y^ could not trigger the decay of a reporter gene containing AU-rich elements (Li et al., 2019). Since the H186Y mutation is located in the second zinc finger region, we tested whether the TZF region is important for TZF1 targeting and subsequent degradation of mRNAs in response to salt stress. RNA immunoprecipitation coupled with qPCR (RIP-qPCR) was performed to determine if any of the down-regulated genes were potential direct targets of TZF1 under salt stress. Eight down-regulated DEGs in *TZF1 OE* plants were chosen from the set of salt stress or oxidative stress related mRNAs containing ARE-like motifs in 3’-UTR (Fig. 6C). Among which, the *autoinhibited Ca^2+^-ATPase* genes *ACA11* and *ACA9* encoding calcium pumps (Li et al., 2023), *KAT3* encoding a subunit of potassium channel (Sun et al., 2015), *ARP1* encoding an ABA-regulated RNA-binding protein 1 (Jung et al., 2013), and *Hsp90.5* encoding a heat shock protein (Song et al., 2009) are all implicated in playing a negative role in salt stress tolerance response. RIP-qPCR results showed that TZF1, but not the TZF^H186Y^, could bind the F1 region of *ACA11* mRNA (Figs. 6D-E). No significant TZF1 binding signals were detected from the rest of the seven potential target genes. These results indicate that *ACA11* downregulation is a potential cause in enhancing salt stress tolerance in *TZF1 OE* plants.

### TZF1 Binds *ACA11* mRNA Directly

To further examine whether TZF1 binds *ACA11* mRNA directly, RNA electrophoretic mobility shift assays (EMSAs) were carried out. The migration of an F4 probe, an extended derivative of the F1 region bound by TZF1 in the RIP-qPCR analysis (Figs. 6D-E), was retarded by GST-TZF1 (Fig. 7A). However, most of the bound complex was unable to enter the gel, so the binding affinity could not be accurately determined. Previous studies showed that both the arginine-rich (RR) and TZF domains of TZF1 were required for high-affinity ARE RNA binding (Qu et al., 2014). In addition, ARE_19_ was shown to be a conserved TZF1 bound RNA motif in 3’-UTR of both plant (Qu et al., 2014) and human (Brooks and Blackshear, 2013) genes. Accordingly, recombinant MBP-TZF1 (RR-TZF) was overexpressed and purified for further EMSAs. Our results show that ARE_19_ shifted readily upon incubation with increasing concentrations of MBP-RR-TZF (Supplemental Fig. S6A). This clear interaction between ARE_19_ and MBP-RR-TZF was leveraged in competition experiments which were used to estimate the binding affinity of ACA11 F1, F4, and F4 deletion derivatives to TZF1. In these assays, unlabeled RNA probes were introduced as competitors to the MBP-RR-TZF–ARE_19_ protein–RNA complex. The resulting gel images showed that F4 could disrupt MBP-RR-TZF binding to ARE_19_ (Fig. 7B). Interestingly, F4 exerted stronger competition than the F1, F4Δ84, and F4Δ134 derivatives (Fig. 7C). Closer examination of the F4 nucleotide sequence revealed two U-rich segments, and each of the deletion derivatives contains only one of these segments. Taken together, these results indicated that TZF1 binds *ACA11* mRNA at the 3’-UTR and that recognition of this region is dependent upon two U-rich segments in the 3’-UTR. To directly assess if TZF1^H186Y^ was defective in mRNA binding, additional EMSAs were conducted. Results showed that MBP-RR-TZF^H186Y^ was unable to bind the consensus ARE_19_ motif (Supplemental Fig. S6A) and that binding between MBP-RR-TZF^H186Y^ and F4 motif was strongly impaired (Supplemental Fig. S6B), suggesting that TZF1^H186Y^ might be defective in mRNA binding in general.

**Figure 7.**
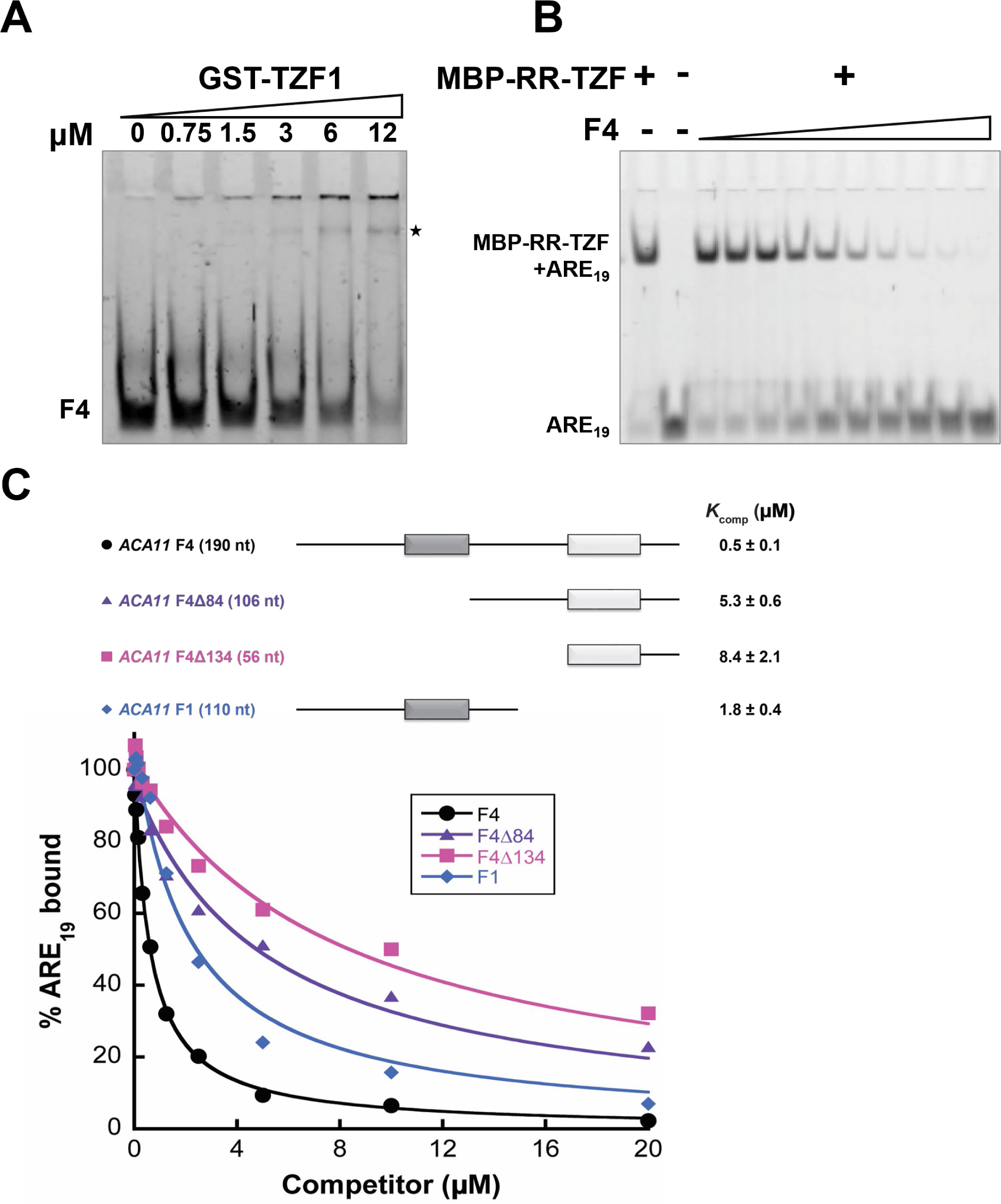
TZF1 binds *ACA11* mRNA in RNA EMSA. A, GST-TZF1 binds the *ACA11* F4 probe region (see Fig. 6D). The RNA-protein complexes are indicated by an asterisk. B, Protein-RNA (MBP-RR-TZF-ARE_19_) complexes are eliminated by competition with unlabeled F4 RNA probe. C, Competition gel shift assays of *ACA11* F1, F4, and F4 deletion derivative RNA probes with ARE_19_ RNA probe. Dark gray and light gray boxes indicate U rich sequence regions.

### TZF1 Enhances the Degradation of *ACA11* mRNA

To investigate if reduced *ACA11* mRNA in *TZF1 OE* lines was due to TZF1 targeting *ACA11* mRNA for degradation, mRNA half-life analysis was conducted. An *Arabidopsis* protoplast transient expression system was used for this assay. In brief, TZF1-mCherry and TZF1^H186Y^-mCherry were used as effectors and GFP-ACA11-F3 (does not bind TZF1) and GFP-ACA11-F4 (binds TZF1) were used as reporters (Fig. 8A). Protoplast samples were co-transformed with a distinct effector and reporter pair and incubated for 10 h before mRNA half-life time-course experiments were conducted using Actinomycin D at 0 h to block the transcription. Results showed that degradation of *GFP*-*ACA11-F4* mRNA was faster in the presence of TZF1 compared to TZF1^H186Y^ (Figs. 8B-C). In contrast, a negligible difference in *GFP*-*ACA11-F3* half-life was observed when it was co-expressed with either TZF1 or TZF1^H186Y^, in agreement with our observation of specific binding of TZF1 to the F1/F4 region of *ACA11* in RIP-qPCR experiments (Figs. 6D-E). To determine if the effectors were expressed at similar levels and the expression of the reporters correlated with corresponding mRNA half-life, we employed fluorescence microscopy. We found that the accumulation of GFP-ACA11-F4, but not GFP-ACA11-F3, was significantly reduced in the presence of TZF1 (Fig. 8D), consistent with the results of mRNA half-life assays (Figs. 8B-C). Immunoblot analysis further confirmed that TZF1 could specifically reduce the accumulation of GFP-ACA11-F4 protein (Fig. 8E). Note that the decay of GFP-ACA11-F4 protein was not obvious after 4 h, likely due to the accumulation and stability of the GFP protein. Together, these results indicate that TZF1 binds and enhances turnover of *ACA11* mRNA.

**Figure 8.**
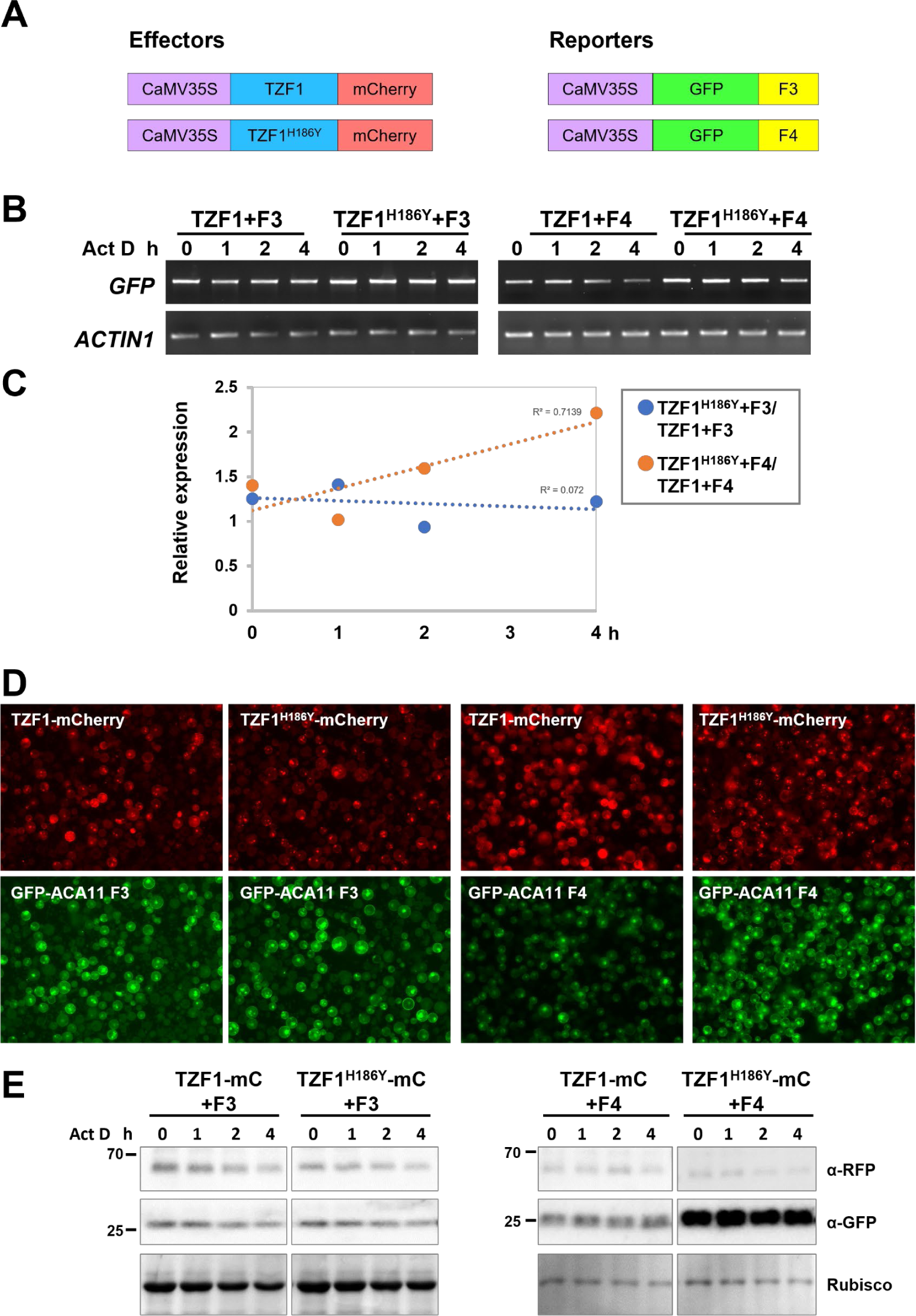
TZF1 triggers turnover of *ACA11-F4* mRNA. A, Constructs used for mRNA half-life analysis. B, Results of RT-PCR analysis indicate that TZF1 can enhance the degradation of *ACA11*-*F4*. Act D: Actinomycin D. C, Quantitative analysis of mRNA half-lives with decay curves/slopes of TZF1^H186Y^+Fn/TZF1+Fn. D, Fluorescence microscopy to determine the protein expression of the effectors and reporters, respectively. The same protoplast sample was imaged with the red channel for mCherry and the green channel for GFP expression. Merged images are not shown. Scale bars = 40 μm. E, Results of the immunoblot analysis indicate that GFP-ACA11-F4 protein is lower when co-expressed with TZF1 than with TZF1^H186Y^. Act D: Actinomycin D. Rubisco protein revealed by Coomassie blue staining was used as a loading control.

### ACA11 Negatively Regulates Salt Stress Tolerance

To determine the role of ACA11 in salt stress tolerance, we compared the salt stress tolerance of wild-type and *aca11* T-DNA insertional mutant under salinity conditions. The *aca11* has the T-DNA inserted into the sixth exon of the *ACA11* (Fig. 9A). RT-PCR results revealed that *ACA11* transcript was absent in homozygous *aca11* line (Figs. 9B-C). In the presence of 200 mM NaCl, *aca11* seedlings showed a significantly higher survival rate than wild-type seedlings (Figs. 9D-E). These results indicated that ACA11 plays a negative role in the salt stress tolerance and TZF1 targeted *ACA11* mRNA for degradation to enhance salt stress tolerance.

**Figure 9.**
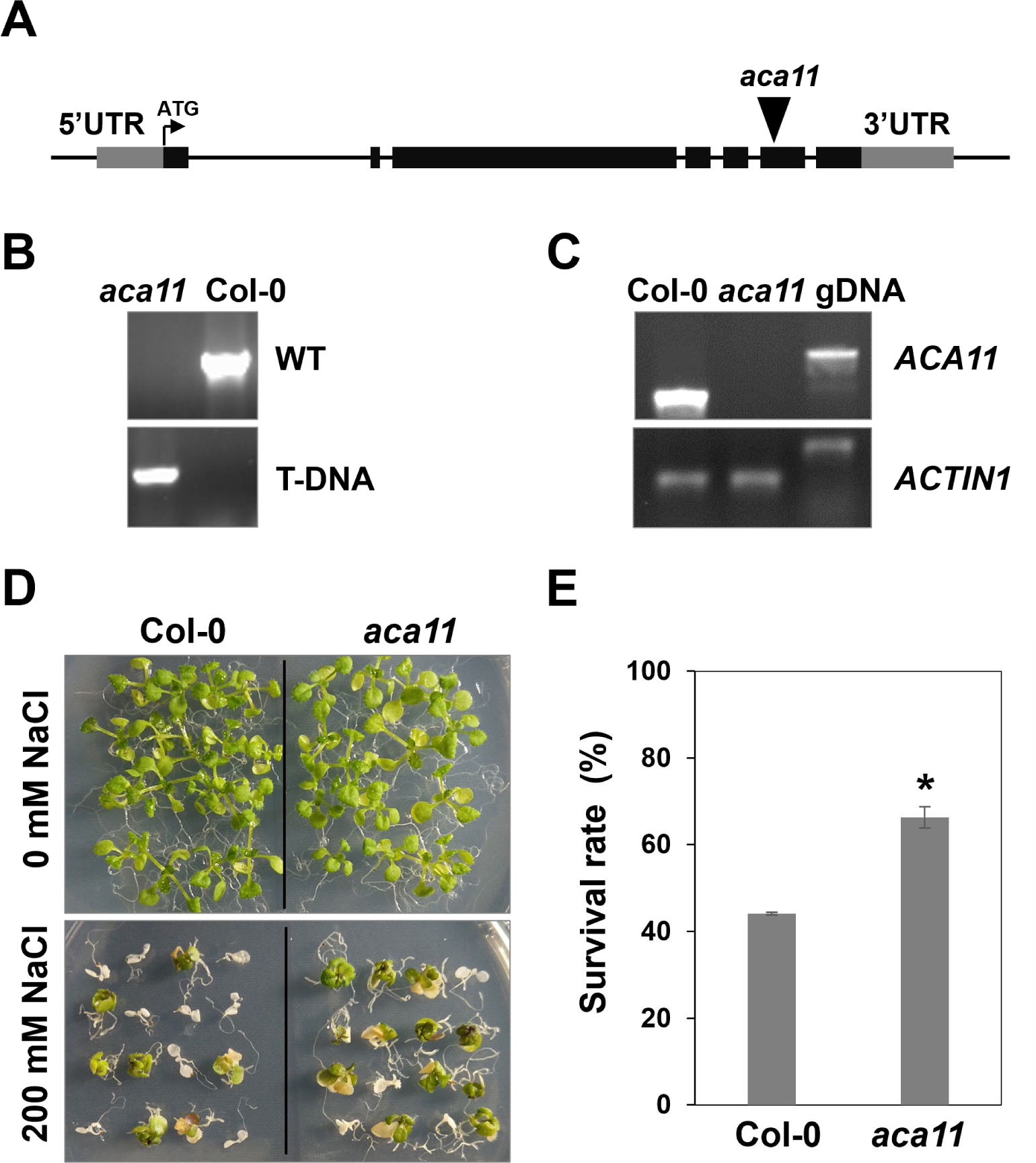
ACA11 acts as a negative regulator of salt stress tolerance. A, Schematic representation of T-DNA insertion in *ACA11* gene. B, PCR genotyping analysis showing that *acall* plant was homozygous for T-DNA insertion. Specific primers were used to detect the wild-type fragment and T-DNA insertion, respectively, in *ACA11* genomic locus. C, Results of RT-PCR analysis indicated that *acall* was a null mutant devoid of *ACA11* transcript. D, The *aca11* seedlings showed enhanced salt stress tolerance than the wild-type plants. Plants were grown on MS plates for seven days and then transferred onto MS medium supplemented with 200 mM NaCl for additional five days. E, Survival rates of seedlings shown in (D). Columns represent means ± s_E_ (n > 20). Asterisks indicate significant differences (*, *P* < 0.05) by Student’s *t* test.

## DISCUSSION

Plant CCCH zinc-finger proteins play pivotal roles in abiotic stress tolerance (Han et al., 2021), but the mechanistic underpinnings are not well understood. In *Arabidopsis*, the expression of *AtTZF1*/*2*/*3*/*10*/*11* were induced by salt treatment, and overexpression of *AtTZF1*/*2*/*3*/*10*/*11* caused enhanced tolerance to salt stress possibly by altered expression of genes involved in biotic/abiotic stress responses (Sun et al., 2007; Lee et al., 2012; Han et al., 2014). Nevertheless, the molecular mechanisms by which AtTZF proteins target mRNAs to control mRNA metabolism in response to salinity stress are still unclear because none of the previous reports showed a complete result of these processes. In this study, the knockout mutant *tzf1* showed a high salt-sensitive phenotype similar to wild-type, while *TZF1 OE* conferred salt stress tolerance in *Arabidopsis* seedlings (Fig. 1; Supplemental Fig. S1). Notably, *tzf1* plants did not show any obvious salt stress-hypersensitive phenotypes, consistent with previous reports of functional redundancy among the TZF family members (Lin et al., 2011). Our data establish unambiguously that TZF1 is a positive regulator of salt stress tolerance in *Arabidopsis*.

Our RNA-seq analysis revealed that many stress-related genes are differentially expressed between wild-type and *TZF1 OE* plants (Figs. 6A-B). A significant number of down-regulated DEGs were related to several abiotic stress-related biological processes including “response to stress” and “response to stimulus”, and down-regulation of such genes is likely to engender the enhanced salt stress tolerance observed in *TZF1 OE* plants. One potential candidate is *ACA11* (Fig. 6C), a tonoplast-localized Ca^2+^ pump (like *ACA4*) (Li et al., 2023). Salt stress is known to result in Ca^2+^ accumulation in the cytoplasm where it functions as an important secondary messenger to trigger a complex signal transduction pathway to enhance salt stress tolerance (Zhao et al., 2021). Countering this flux, ACA11 and ACA4 expel Ca^2+^ from the cytoplasm to vacuoles, thereby acting as negative regulators for Ca^2+^ accumulation and dampening the salt-stress protective response. Consistent with this function, the double-knockout *aca4*/*aca11* leaves exhibit elevated baseline cytoplasmic Ca^2+^ levels (Hilleary et al., 2020). The transcript level of *ACA11* was reduced in *TZF1 OE* (Fig. 6), and both *TZF1 OE* plants and *aca11* mutants with reduced *ACA11* expression displayed an enhanced salt stress tolerance phenotype (Figs. 1 and 9). These results indicate that TZF1 positively regulates salt stress tolerance, partly by down-regulating *ACA11* through mRNA binding and degradation in *Arabidopsis*. To determine whether TZF1 is involved in mRNA turnover of target genes, we searched for ARE motifs in the down-regulated genes in *TZF1 OE* plants under salinity stress. Although we found eight genes containing typical ARE/URE motifs in their 3’-UTR (Fig. 6C), we focused our attention on *ACA11*. Our results showed that AtTZF1 could bind *ACA11* mRNA in specific U-rich regions in 3’-UTR both *in vivo* and *in vitro* (Figs. 6E and 7). We further demonstrated that TZF1 binds both ARE and *ACA11* 3’-UTR with higher affinities than TZF1^H186Y^ (Supplemental Fig. S6). Using mRNA half-life analysis, we established a correlation between TZF1 binding and enhanced mRNA degradation. Interestingly, we found that *ACA4* was also strongly downregulated by salt stress in *TZF1 OE* but not *TZF1^H186Y^ OE* plants in the RNA-seq analysis (Supplemental Fig. S7). Although *ACA4* does not contain typical AREs or UREs at 3’-UTR, we cannot rule out the possibility that TZF1 could also bind *ACA4* and trigger mRNA degradation. Together, these results indicate that *TZF1 OE* plants engender decreased expression of *ACA11* and *ACA4*, and thereby confer salt stress tolerance presumably by allowing a longer duration of elevated cytoplasmic Ca^2+^ levels (Fig.10), a pre-requisite for triggering the protective response to salt stress.

**Figure 10.**
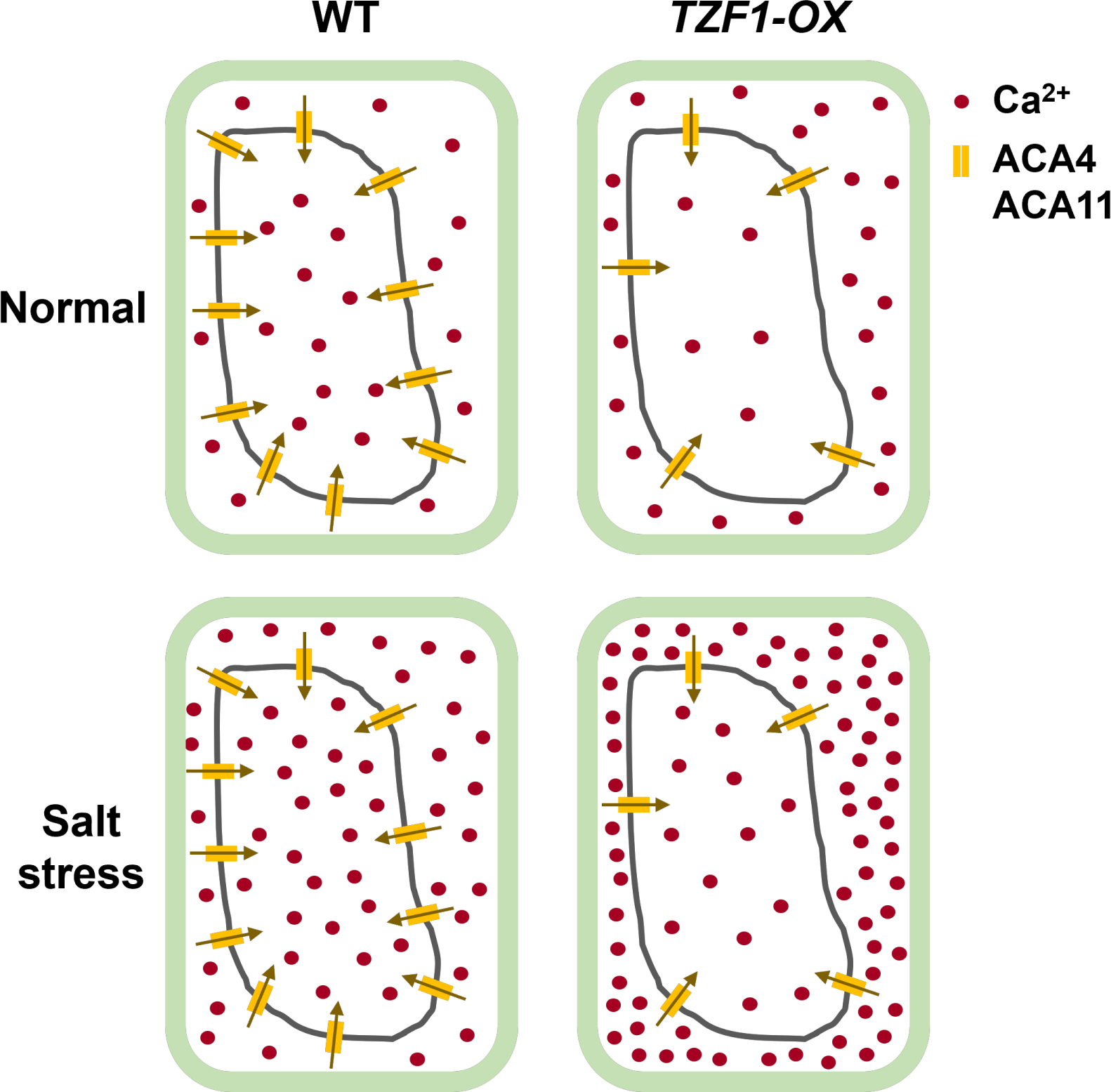
Schematic model of increased Ca^2+^ distribution in the cytoplasm under salt stress in *TZF1 OE* plants. Salt stress induces cytoplasmic Ca^2+^ accumulation. ACA4 and ACA11 are two major tonoplast-localized calcium pumps that expel Ca^2+^ from cytoplasm into the large central vacuole during salt stress. In *TZF1 OE* plants, decreased expression of *ACA4* and *ACA11* hinders the calcium transport into the vacuole hence maintaining high Ca^2+^ concentration in the cytoplasm, supporting an enhanced salt stress tolerance response.

Our results also inspire many future investigation avenues that are likely to be profitable. *First*, like *ACAs*, *KAT3* (*AtKC1*), a potassium channel protein, was also downregulated in *TZF1 OE* plants (Fig. 6C). As mentioned earlier, the low cytosolic Na^+^/K^+^ ratio is a key indication of salt stress tolerance (Zhao et al., 2021). One of the natural mechanisms of enhancing salt stress tolerance is to retain high K^+^ level in the cytoplasm (Sun et al., 2015). KAT3 acts as a regulatory subunit within the heterotetrameric (KAT3, KAT1, KAT2, and AKT2) potassium channel, reducing the K^+^conductance and negatively shifting the potassium channel activation potential (Jeanguenin et al., 2011). Therefore, downregulation of *KAT3* in *TZF1 OE* plants could potentially activate the potassium channel and maintain high cytoplasmic K^+^ level to enhance salt stress tolerance. Other than *ACAs* and *KAT3*, the downregulation of *ARP1* and *Hsp90.5* (Fig. 6C) might also contribute to salt stress tolerance in *TZF1 OE* plants. Aberrant expression of *ARP1* delayed seed germination under ABA, high salt, or dehydration stress conditions (Jung et al., 2013), while overexpression of *AtHsp90.5* reduced tolerance to both salt and drought stresses (Song et al., 2009). Additional experimentation is necessary to parse the individual contributions of these different players.

*Second*, based on the inventory of up-regulated DEGs which include several positive regulators of plant salt stress tolerance (Supplemental Fig. S5C), we hypothesize that TZF1 might target the degradation of mRNAs that encode repressors of these positive regulators. For example, the *NHX* gene family contributes to Na^+^ homeostasis in plants and plays an important role in conferring salinity tolerance (Fang et al., 2021). *NHX1* encodes a vacuolar sodium/proton antiporter involved in salt tolerance and ion homeostasis. Overexpression of *NHX1* improved salt tolerance in transgenic plants of several species (Apse et al., 1999; Zhang and Blumwald, 2001; Apse and Blumwald, 2007; Kumar et al., 2017). *NHX2* encodes a vacuolar K^+^/H^+^ exchanger essential for active K^+^ uptake at the tonoplast. Overexpression of Jerusalem artichoke *NHX2* in rice improved salt stress tolerance (Zeng et al., 2018). *NHX6* encodes an endosomal Na^+^/H^+^ antiporter, double knockout *nhx5nhx6* showed increased sensitivity to salinity (Bassil et al., 2011). For the rest of the genes listed in Supplemental Fig. S5C, the *AtPP2-B11* plays an important role in response to salt stress by up-regulating *AnnAt1* expression, repressing ROS production, and disrupting Na^+^ homeostasis in *Arabidopsis* (Jia et al., 2015). Under salt stress, *VPS23A* positively regulates the redistribution of SOS2 protein, also known as CIPK24, to the plasma membrane, which then activates the Na^+^/H^+^ antiporter SOS1 (known as NHX7) activity to exclude excess cytoplasmic Na^+^ and confers salt stress tolerance in plants (Lou et al., 2020). Salt-induced stabilization of RGL3 by nitric oxide (NO) confers enhanced salt stress resistance (Shi et al., 2017). *CIPK8* is involved in regulating plant salt tolerance by promoting Na^+^ export from cells (Yin et al., 2020).

*Last*, a more complete cell biological study on TZF-mRNA interaction and target mRNA metabolism is critical. Plant TZF proteins can localize to PBs and SGs and bind specific RNA elements to trigger RNA degradation. However, very few *in vivo* mRNA binding targets have been identified to date. AtTZF1 was shown to localize in PBs and SGs and could trigger ARE-containing mRNA degradation *in vivo* (Pomeranz et al., 2010; Qu et al., 2014). In addition, AtTZF1 negatively regulates TOR signaling by binding to the 3’-UTR and promoting *TOR* mRNA degradation (Li et al., 2019). OsTZF1 and OsTZF7 can bind mRNAs of down-regulated genes containing U-rich and ARE-like motifs within their 3’-UTRs (Jan et al., 2013; Guo et al., 2022). Cotton GhTZF2 is localized in cytoplasmic granules and the transcriptome analysis showed that many differentially expressed transcripts were enriched with AREs in their 3’-UTRs, suggesting that GhTZF2 might regulate mRNA turnover of target genes (Li et al., 2023). Tomato TZF protein SIC3H39 could colocalize with PB marker DCP2 and SG marker PABP8. It negatively regulated cold stress tolerance by binding to the 3’-UTR AREs and triggering degradation of the cold-responsive mRNAs (Xu et al., 2023). Despite numerous studies consistently implying that TZF proteins enhance abiotic stress response by degrading target mRNAs, the molecular details between the steps of mRNA targeting and degradation are missing. In this report, we have unequivocally determined that TZF1 protein is mainly localized in SGs (Fig. 4). Whether or not the degradation of target mRNAs takes place in SGs, PBs, or both remains an important question to address in the future. Nevertheless, we have demonstrated that both TZF1 and TZF1^H186Y^ accumulated in SGs under salinity stress (Fig. 3; Supplemental Fig. S4), suggesting that the localization of TZF1 to cytoplasmic granules might not be a sole prerequisite for its function in regulating the fates of target mRNAs. The subcellular localization and function of TZF1 in salt stress tolerance might be regulated through different mechanisms. Through deletion and site-directed mutagenesis analyses, we found that TZF1 localization to SGs is controlled by multiple domains and numerous post-translational modification mechanisms (data not shown). For example, TZF1 protein without TZF domain could still be localized to SGs, suggesting that the mechanisms governing subcellular localization and RNA binding and/or decay could be uncoupled, given both this and previous reports (Qu et al., 2014) indicating that the RR-TZF motif is responsible for RNA-binding. By contrast, deletion of IDRs from TZF1 significantly reduced its assembly into SGs. Whether or not these IDR deletion TZF1 mutants could still confer salt stress tolerance remains an outstanding future task.

In retrospect, the higher abundance of cytoplasmic granules observed in *TZF1^H186Y^ OE* plants could be due to the following possibilities: *1*) TZF1^H186Y^-GFP protein is more stable than TZF1-GFP protein (Figs. 3 and 5; Supplemental Fig. S4). It is remarkable that a single amino acid change could change TZF1 from being extremely labile to relatively stable (Fig. 5B). *2*) TZF1^H186Y^ does not bind *ACA11* but could still bind other mRNAs and be sequestered into cytoplasmic granules. *3*) TZF1^H186Y^ could interact with additional mRNA binding proteins and hence be sequestered into cytoplasmic granules. Conversely, TZF1 cytoplasmic granule localization is indeed coupled with its function in target mRNA degradation. An alternative hypothesis for this possibility is that *ACA11* mRNA degradation by TZF1 might mainly occur in the cytoplasm, and stronger sequestration of TZF1^H186Y^ to cytoplasmic granules might prevent its accessibility to the *ACA11* mRNA in the cytoplasm. Together with lower ability to bind and promote degradation of *ACA11* mRNA, *ACA11* mRNA hence accumulates at a higher level in *TZF1^H186Y^ OE* plants. Details of the interplay between subcellular localization and mRNA turnover remain to be worked out in the future studies.

In sum, we propose that TZF1 exerts a surprisingly effective, two-pronged post-transcriptional regulatory control on the expression of a specific set of genes involved in salt stress tolerance. By downregulating the expression of vacuolar Ca^2+^ pumps (*ACA11* and *ACA4*), the likelihood of salt stress alleviation and a cellular homeostasis reset is enhanced (Fig. 10). Although we lack the identity of the repressors targeted by TZF1, a second front to offset the sodium assault is its indirect contribution to increased expression of Na^+^/H^+^ exchangers and thereby provide an off-ramp to dampen rising cytosolic Na^+^ levels. Moving forward, this regulatory axis of TZF1-Ca^2+^-salt stress response could be leveraged to improve crop salt stress tolerance. For example, plants expressing a salt-inducible promoter fused with *TZF1* gene could enhance salt stress tolerance but bypassing the developmental abnormalities caused by the constitutive expression of *TZF1* driven by a ubiquitous promoter such as *CaMV35S* (Lin et al., 2011).

## MATERIALS AND METHODS

### Plant Materials and Growth Conditions

Wild-type *Arabidopsis thaliana* Columbia-0 (Col-0), *TZF1 OE*, *TZF1^H186Y^OE*, *tzf1* (SALK_143721), and *aca11* (SALK_060930) plants were used in this study. The plant seeds were sterilized and plated on Murashige and Skoog (MS) medium (Phytotech) for 2 days at 4°C, and then transferred to a growth chamber with 16/8 h light/dark cycles at 22°C. The light intensity was 50 µmol m^−1^ s^−1^.

### Chlorophyll Content Measurement

Sixteen seedlings were placed in a 2 ml centrifuge tube and the plant fresh weight was recorded as W. To each tube, 1 ml aliquots of 80% (v/v) acetone were added to immerse the plants. The samples were then placed in the dark for 48 h at 22℃. The acetone extract was isolated and its volume was recorded as V. For each extract, absorbance at 645 nm and 663 nm was measured using a spectrophotometer (TECAN Infinite M200 Pro) and chlorophyll content was calculated using the formula [(8.02 × *Abs*_663_) +(20.21 × *Abs*_645_)]×V/W as previously described (Li et al., 2021).

### Na^+^ and K^+^ Content Measurement

Accumulation of Na^+^ and K^+^ ions was measured according to a previously described (Jiang et al., 2019). Plants grown in the presence or absence of NaCl treatment were separately harvested and dried for 48 h at 65℃ before their weight was measured. All samples were then digested with 75% (v/v) nitric acid and 25% (v/v) hydrogen peroxide at 180°C for 3 h. The samples were then diluted to 30 ml with ddH_2_O and filtered. The Na^+^ and K^+^ contents in each solution was measured using an inductively coupled plasma optical emission spectrometer (ICAP6300).

### DAB Staining

Seedlings and leaves were incubated in DAB staining solution (1 mg/ml DAB dissolved in ddH_2_O, pH 3.8) in the dark at 22℃ for 8 h. The samples were then destained with acetic acid:glycerin:ethanol (1:1:4 by volume) at 80℃ for 25 min before imaging.

### RNA-seq Sample Preparation and Sequencing

Eleven-day-old seedlings grown on MS plates under 12/12 h light/dark cycles were treated with 150 mM NaCl at Zeitgeber time ZT2 for 4 h before sample collection and total RNA extraction. Three biological replicates were prepared for each sample. Total RNA was extracted using Trizol reagent (Invitrogen). The RNA was then treated with DNase I (DNA-*free*™ DNA Removal Kit, Invitrogen) to remove genomic DNA. RNA quality was evaluated on a Bioanalyzer 2100 instrument (Agilent). Sequencing libraries were prepared using the Directional RNA Library Prep Kit (E7760S, New England Biolabs) according to manufacturer’s instructions. The 150-nucleotide (nt) paired-end high-throughput sequencing was performed using an Illumina Hiseq X TEN platform. After removing the low-quality sequencing reads, the remaining clean reads were mapped to the *Arabidopsis* reference genome (TAIR10) using Tophat2 software. DEGs were analyzed using edgeR software. Genes with *q* < 0.05 and |log 2 _ratio| ≥ 1.5 were identified as DEGs. The enrichment of DEGs in different functional categories was performed by using agriGO (GeneOntology) V2.0. Venn diagram analysis was conducted using VENNY 2.1. Three biological replicates were prepared for each sample.

### RNA Immunoprecipitation (RIP) and RT-qPCR Analysis

Eleven-day-old seedlings grown on MS plates under 12/12-h light/dark cycles were treated with 150 mM NaCl at ZT2 for 4 h before samples were collected for crosslinking. RIP was performed as previously described (Koster et al., 2014). Briefly, the cross-linked tissues (1.5 g) were ground with liquid nitrogen and resuspended in 750 µl prewarmed (60℃) RIP lysis buffer to make viscous homogenates. After centrifugation, the cell extract supernatant was filtered through a 0.45 µm filter. The extract was then pre-cleared twice with 50 µl Sepharose beads before mixing with 15 µl washed Sepharose beads coated with GFP antibody (Invitrogen). The beads were incubated with extract for 2 h at 4℃ and then washed three times with RIP washing buffer for 10 min at 4℃. The beads were then washed with RIP lysis buffer for 5 min at 4℃. The RNA was purified from the immunoprecipitated RNPs and 100 µl input, respectively, using the Trizol reagent (Invitrogen). For quantitative analysis of TZF1 binding to RNA, reverse transcription was performed using SuperScript™ IV First Strand Synthesis System. After DNase treatment, RNA samples were reverse transcribed with random hexamer primers. Quantitative PCR was performed to determine the level of TZF1-bound RNA, and the 2^−ΔCT^ method was used to calculate the ratio of RIP to the input.

### TZF1 Protein Half-Life Analysis

Seven-day-old *TZF1 OE* and *TZF1^H186Y^ OE* seedlings were incubated with 30 μM cycloheximide (CHX), 50 μM MG115/132, or both CHX and MG115/132. Seedlings were collected at different time points, and then total proteins were extracted and protein levels were determined by immunoblot analysis with anti-GFP antibody (Roche).

### Recombinant Protein Production

Recombinant GST-TZF1 and MBP-TZF1 (RR-TZF) proteins were produced by using *Escherichia coli* BL21 (DE3) cells. Bacterial cultures were grown to *Abs*_600_ = 0.6, at which point 0.1 mM isopropyl β-D-1-thiogalactopyranoside (IPTG) was added for induction of protein production. ZnCl_2_ (0.1 mM) and glucose [0.2% (w/v); MBP-RR-TZF proteins only] were also added at the time of addition induction. Cultures were then grown for an additional 12-16 h at 18°C before harvesting. GST-TZF1 was purified using glutathione-Sepharose 4B resin (GE Healthcare Bio Sciences) as previously described (Qu et al., 2014). For purification of MBP-RR-TZF, cell pellets from a 1 L overexpression culture were resuspended in 20 ml extraction buffer [50 mM Tris-HCl, pH 8.5; 100 mM NaCl; 1 mM EDTA, 1 mM DTT, 1 mM PMSF, 0.5X EDTA-free protease inhibitor cocktail (APExBIO)]. The resuspended cells were lysed at 15,000 psi using a French pressure cell press, and the resulting whole-cell lysate was supplemented with 0.2% (v/v) Triton X-100 and incubated at 4°C for 10 min with gentle nutation. The lysate was then supplemented with 10 mM MgCl_2_ and 10 mM CaCl_2_ and 5 µg DNase I (Sigma-Aldrich) and incubated at 20°C for an additional 10 min. After incubation, the lysate was clarified by centrifugation at 12,000 x *g* and 4°C for 15 min and then filtered using a 0.45-µm syringe filter (Roche). The filtered lysate was passed over a gravity-flow column packed with 0.5 ml amylose resin (New England Biolabs) that had been pre-equilibrated with extraction buffer. After collecting the flowthrough, the amylose resin was washed with 20 ml extraction buffer to remove weakly bound proteins. Bound MBP-RR-TZF was eluted with 0.5 ml additions of elution buffer (50 mM Tris-HCl, pH 8.5, 10 mM amylose, 1 mM DTT) and individual elution fractions were collected and assessed by SDS-PAGE. The fractions containing near-homogeneous MBP-RR-TZF were pooled and dialyzed twice against 500 ml storage buffer (20 mM Tris-HCl, pH 8.0; 50 mM NaCl; 1 mM DTT) at 4°C. The final protein concentration was determined by measuring *Abs*_280_ (ε = 81,820 M^-1^cm^-1^) and aliquots were stored at -80°C. Prior to use in competition assays, aliquots of MBP-RR-TZF were thawed and concentrated to ∼ 80 µM using a Vivaspin 500 10,000-MWCO centrifugal concentrator (Sartorius). Concentrated protein aliquots were stored at -80°C.

### Preparation of RNAs

The F4, F4Δ84, F4Δ134, and F1 RNAs were generated by run-off *in vitro* transcription (IVT) using T7 RNA polymerase. The DNA templates for IVT were prepared by PCR using cDNA as the template. The F4, F4Δ84, and F4Δ134 IVT templates were amplified using F4-F, F4Δ84-F, and F4Δ134-F forward primers, respectively, and F4-R as the common reverse primer. Due to poor amplification of the F4Δ134 template after the first PCR, the amplicon from the first round was used as the template in a second PCR that used the same set of primers as in the first round. IVT reactions contained 1x IVT buffer [40 mM Tris-HCl, pH 7.6; 24 mM MgCl_2_; 2 mM spermidine; 0.01% (v/v) Triton X-100]; 10 mM DTT; 5 mM each of ATP, CTP, GTP, and UTP; 0.002 U thermostable inorganic pyrophosphatase (NEB); 0.15-0.3 µg template, and T7 RNA polymerase (purified in-house)]. After 4 h at 37°C, the reactions were treated with DNase I (Roche), extracted with phenol-chloroform, and dialyzed using 3,500-MWCO tubing (BioDesign Inc. New York) against 3.5 L ddH_2_O three times over 20 h at 4°C and once over 1.5 h at 20°C. Following dialysis, the RNAs were precipitated with 0.3 M sodium acetate and 2.5 volumes of ethanol, resuspended in autoclaved ddH_2_O, and quantitated by measuring the absorbance at *Abs*_260_ and using their respective extinction coefficients (OligoAnalyzer, IDT).

To prepare 5′-[^32^P]-labeled F4 (5′-[^32^P]-F4), F4 was first dephosphorylated by incubating with calf intestinal phosphatase (NEB) for 2 h at 37°C. Dephosphorylated F4 was then extracted with phenol-chloroform, precipitated with sodium acetate and ethanol, and resuspended in ddH_2_O as described above. Following clean-up, dephosphorylated F4 (5 µM) was incubated with γ-[^32^P]-ATP and T4 polynucleotide kinase (NEB) for 45 min at 37°C. The labeling reaction was quenched with the addition of urea dye [7 M urea, 1 mM EDTA, 0.05% (w/v) xylene cyanol, 0.05% (w/v) bromophenol blue, 10% (v/v) phenol], loaded onto a denaturing 8% (w/v) polyacrylamide/7 M urea/1X Tris-Borate-EDTA (TBE) gel, and electrophoresed in 1X TBE buffer for 75 min. Full-length 5′-[^32^P]-F4 was identified in the gel by autoradiography, excised from the gel, and eluted using the crush-and-soak method. Eluted 5′-[^32^P]-F4 was precipitated with sodium acetate and ethanol and resuspended to a final specific activity of 200,000 dpm/µl in ddH_2_O.

### Gel-shift Assays

For TZF binding to ARE_19,_ a pre-mixed solution of 100 nM 6-FAM-labeled ARE_19_ (6-FAM-ARE_19_, IDT) and 40 µM 20-nt decoy oligo (MilliporeSigma) in binding buffer [20 mM HEPES-KOH, pH 7.5; 50 mM NaCl; 5 µM ZnCl_2_, 4 mM MgCl_2_] was aliquoted into multiple tubes. Importantly, inclusion of the 20 nt decoy oligo minimized apparent nucleolytic decay of the target RNA during the binding reaction and improved the resolution of free 6-FAM-ARE_19_ bands in the gel image. MBP-RR-TZF (or H186Y mutant) was serially diluted from 40 μM to 156 nM by diluting with equal volumes of storage buffer [20 mM Tris-HCl, pH 8; 50 mM NaCl; 1 mM DTT]. To initiate the binding reaction, equal volumes of MBP-RR-TZF were combined with the pre-mixed aliquots of 6-FAM-ARE_19_ and decoy oligo. As a negative control, a single aliquot of 6-FAM-ARE_19_ and decoy oligo was diluted with an equal volume of storage buffer. The binding reactions were incubated at 23°C for 10 min. Loading dye containing xylene cyanol, bromophenol blue, and 50% (v/v) glycerol was added to each tube, and each reaction mix was loaded onto a native 8% (w/v) polyacrylamide (29:1 acrylamide:bisacrylamide)/1X Tris-Borate (TB) gel. The gel was electrophoresed in 1X TB buffer at 100 V for 1 h at 4°C and then scanned using an Amersham Typhoon Biomolecular Imager (Cytiva) on the Cy2 setting.

For TZF binding to ACA11-F4, a pre-mixed solution of 1 nM 5′-[^32^P]-F4 and 40 µM 20-nt decoy oligo in binding buffer was aliquoted into multiple tubes. MBP-RR-TZF (or H186Y mutant) was serially diluted from 40 μM to 156 nM by diluting with equal volumes of storage buffer. To initiate the binding reaction, equal volumes of MBP-RR-TZF were combined with the pre-mixed aliquots of 5′-[^32^P]-F4 and decoy oligo. The binding reactions were incubated and resolved by native polyacrylamide gel electrophoresis as described for 6-FAM-ARE_19_, except the binding reactions were resolved on a 6% (w/v) polyacrylamide/1X TB gel, and the gel was exposed to storage phosphor screen which was then scanned using an Amersham Typhoon Biomolecular Imager (Cytiva) on the phosphorimager setting.

### Competition Gel Shift Assays

The binding affinity of MBP-RR-TZF for ACA11 F4 and deletion derivatives (F1, F4Δ84, F4Δ134) was measured in a gel-shift assay by monitoring dissociation of MBP-RR-TZF from 6-FAM-ARE_19_, which was previously established as a minimal RNA ligand for MBP-RR-TZF (Qu et al., 2014). For all competition assays, a fixed concentration of 20 µM MBP-RR-TZF was used to ensure near-complete binding to ARE_19_ in the absence of competitor RNA, and then subject to a competition with increasing concentration of either F4 or one of the F4 deletion derivatives. All reported RNA and protein concentrations reflect the final concentrations in the binding assay. A pre-mixed solution of 50 nM 6-FAM-ARE_19_ and 20 µM 20-nt decoy oligo in binding buffer (20 mM HEPES-KOH, pH 7.5; 50 mM NaCl; 5 µM ZnCl_2_, 4 mM MgCl_2_) was aliquoted into multiple tubes and supplemented with increasing concentrations of competitor RNA. Two aliquots that were designated as positive- and negative-binding controls were supplemented with ddH_2_O instead of competitor RNA. To the binding reaction and positive-control tubes, 20 µM MBP-RR-TZF was added, whereas storage buffer was added to the negative control tube. The binding reactions were incubated and resolved by native polyacrylamide gel electrophoresis as described for 6-FAM-ARE_19_. Following electrophoresis, gels were scanned using an Amersham Typhoon imager (Cytiva) on the Cy2 setting, and the intensity of bands corresponding to bound and unbound 6-FAM-ARE_19_ were quantified using ImageQuant software (Molecular Dynamics). The fraction of 6-FAM-ARE_19_ bound at each competitor concentration was normalized using the fraction of ARE_19_ bound in the positive control sample and plotted as % bound ARE_19_. Data were fit to a hyperbola ([ΔARE_19_ bound]\**K_comp_*)/([competitor] + *K*_comp_) to yield the *K*_comp_ value for binding of MBP-RR-TZF to a competitor RNA. All reported *K*_comp_ values and errors represent average *K*_comp_ and standard deviation determined from three technical replicates.

### Messenger RNA half-life Assays

The TZF1, TZF1^H186Y^, ACA11-F3, and ACA11-F4 were cloned into the pENTR^™^/D-TOPO^®^ vectors. All constructs were subcloned into the Gateway^®^ destination vectors with N-terminal GFP or C-terminal mCherry tag by using the LR recombination reaction. *Arabidopsis* protoplast samples were co-transformed with a distinct effector and reporter pair (TZF1+ACA11-F3, TZF1+ACA11-F4, TZF1^H186Y^+ACA11-F3, TZF1^H186Y^+ACA11-F4) and incubated for 10 h before mRNA half-life time-course experiments were conducted. Actinomycin D at 100 μg/ml was used (Sigma) to block the transcription at the beginning of the time-course experiments. Total RNA was extracted by using a RNeasy plant mini kit (Qiagen), then treated with TURBO DNA-*free*^TM^ Kit (Ambion) to remove contaminating genomic and plasmid DNA. First-strand cDNA was synthesized by using the SuperScript III reverse transcriptase (Invitrogen) according to the supplier’s instructions. *ACTIN1* was used as an internal control for RT-PCR analysis.

### Accession Numbers

Sequence data from this article can be found in the Arabidopsis Genome Initiative under the following accession numbers: TZF1 (AT2G25900) and ACA11 (AT3G57330).

## SUPPLEMENTAL DATA

**Supplemental Figure S1.** The T-DNA knockout mutant *tzf1* showed salt stress-sensitive phenotypes.

A, Phenotypes of the Col-0, *TZF1 OE*, *TZF1^H186Y^ OE*, and *tzf1* plants under normal and salt stress conditions. Seven-day-old seedlings grown under long-day condition (16/8 h light/dark cycles) were transferred to MS plates containing different NaCl concentrations for eight additional days. B, Survival rates of seedlings shown in (A). Data represent the average of 3 replicates ± *SD*. Different letters (*a* and *b*) indicate significant differences at *P*< 0.05 by one-way ANOVA analysis using the SPSS software.

**Supplemental Figure S2.** TZF1 is not involved in the osmotic stress response induced by sorbitol.

Phenotypes of the Col-0, *TZF1 OE*, *TZF1^H186Y^ OE*, and *tzf1* plants under osmotic stress induced by sorbitol. Seven-day-old seedlings grown under long-day condition (16/8 h light/dark cycles) were transferred to MS plates containing different concentrations of sorbitol for eight additional days.

**Supplemental Figure S3.** Recapitulation of salt stress sensitive phenotypes of EMS allele of *TZF1^H186Y^ OE* plants by ectopic expression of *TZF1^H186Y^-GFP* in the wild-type background (*TZF1^H186Y^ OE L8*).

A, Phenotypes of the Col-0, *TZF1 OE* (*L83*), and *TZF1^H186Y^ OE* (*L8*) plants under normal and salt stress conditions. Seven-day-old seedlings grown under long-day condition (16/8 h light/dark cycles) were transferred to MS plates containing different NaCl concentrations for additional eight days. B, Survival rates of seedlings shown in (A). Data represent the average of 3 replicates ± *SD*. Different letters (*a* and *b*) indicate significant differences at *P*< 0.05 by one-way ANOVA analysis using the SPSS software.

**Supplemental Figure S4.** TZF1 cytoplasmic foci induced by salt stress.

Fluorescent micrographs of root cells of seven-day-old *TZF1 OE* and *TZF1^H186Y^ OE* etiolated seedlings treated with 200 mM NaCl for 4 h. Consistent with the results shown in Figure 3, salt-induced cytoplasmic foci appeared to be more abundant in *TZF1^H186Y^ OE* plants.

**Supplemental Figure S5.** RNA-seq analysis reveals up-regulated DEGs in responses to salt stress.

A, Venn diagram analysis of up-regulated DEGs in the Col-0, *TZF1 OE*, and *TZF1^H186Y^ OE* plants treated with NaCl. RNA-seq analysis was conducted using eleven-day-old seedlings grown on MS plates under 12 h light/dark cycles and treated with 150 mM NaCl for 4 h. B, GO analysis of up-regulated DEGs specifically found in NaCl-treated *TZF1 OE* plants. *P* values were generated by Fisher’s exact test. C, A subset of up-regulated DEGs encoding positive regulators of salt stress tolerance found in NaCl-treated *TZF1 OE* plants.

**Supplemental Figure S6.** The RR-TZF fragment of TZF1 protein binds to ARE_19_ and ACA11 F4 RNA probes in EMSA.

A, The recombinant MBP-RR-TZF protein binds to ARE_19_ in EMSA. MBP-RR-TZF or MBP-RR-TZF^H186Y^ mutant (78 nM to 20 μM final) was incubated with 50 nM 6-FAM-ARE_19_ at 23°C for 10 min, and complex formation was subsequently analyzed by native 8% (w/v) polyacrylamide gel electrophoresis. The RNA-protein complexes are indicated by an asterisk. B, The recombinant MBP-RR-TZF protein binds to ACA11 F4 in EMSA. MBP-RR-TZF or MBP-RR-TZF^H186Y^ mutant (78 nM to 20 μM final) was incubated with 0.5 nM 5′-[^32^P]-F4 RNA at 23°C for 10 min, and complex formation was subsequently analyzed by native 6% (w/v) polyacrylamide gel electrophoresis. The intensity of the shifted band was higher for the MBP-RR-TZF protein than for the MBP-RR-TZF^H186Y^, indicating tighter binding of F4 by the RR-TZF protein. The RNA-protein complexes are indicated by asterisks.

**Supplemental Figure S7.** *ACA4*, *ACA9*, and *ACA11* are downregulated in *TZF1 OE* plants under salt stress.

Normalized relative expression of *ACA4*, *ACA9*, and *ACA11* was determined from 3 biological RNA-seq analysis replicates.

**Supplemental Table S1.** DNA oligonucleotides used in this study.

## AUTHOR CONTRIBUTIONS

S.-L.H., B.L., W.J.Z., V.G., L.W., and J.-C.J. conceived and designed the experiments; S.-L.H. B.L., W.J.Z., H.C.A., and V.S. performed the experiments; S.-L.H. and J.-C.J. wrote the manuscript with input from all authors.

## ACKNOWLEDGMENTS

We thank Ms. Jingquan Li from the Key Laboratory of Plant Molecular Physiology and Plant Science Facility of the Institute of Botany, CAS for their technical assistance of confocal microscopy assay.

